# Overexpression of *βTrCP1* elicits cell death in cisplatin-induced senescent cells

**DOI:** 10.1101/2024.11.12.622981

**Authors:** Alejandro Belmonte-Fernández, Joaquín Herrero-Ruíz, M. Cristina Limón-Mortés, Carmen Sáez, Miguel Á. Japón, Mar Mora-Santos, Francisco Romero

## Abstract

Senescence is a non-proliferative cellular state derived from aging or in response to exogenous insults, such as those that cause DNA damage. As a result of cancer treatments like cisplatin, certain tumor cells may undergo senescence. However, rather than being beneficial for patients, this is detrimental because these cells might proliferate again under specific conditions and, more importantly, because they synthesize and secrete molecules that promote the proliferation of nearby cells. Therefore, to achieve complete tumor remission, it is necessary to develop senolytic compounds to eliminate senescent cells. Here, we studied the role of βTrCP1 in cell proliferation and senescence and found that lentiviral overexpression of *βTrCP1* induces the death of senescent cells obtained after cisplatin treatment in both two-dimensional cell cultures and tumorspheres. Mechanistically, we demonstrated that overexpression of *βTrCP1* triggers proteasome-dependent degradation of p21 CIP1, allowing damaged cells to progress through the cell cycle and consequently die. Furthermore, we identified nucleophosmin 1 (NPM1) as the intermediary molecule involved in the effect of βTrCP1 on p21 CIP1. We determined that increased amounts of βTrCP1 partially retains NPM1 in the nucleoli, preventing it from associating with p21 CIP1, thus leaving it unprotected from degradation by the proteasome. These results have allowed us to discover a potential new target for senolytic drugs, as retaining NPM1 in the nucleoli under senescent conditions induces cell death.

## INTRODUCTION

Beta-transducin repeat-containing protein (βTrCP) is an F-box protein that forms part of the SCF (SKP1-CUL1-F-box protein) E3 ubiquitin ligase complex. These complexes participate in the ubiquitylation of proteins involved in a variety of cellular processes, including the cell cycle, apoptosis or metabolism ^1^. Specifically, the F-box proteins are responsible for recognizing substrates to be ubiquitylated by the enzymatic machinery. These ubiquitylated proteins can follow various pathways: they can be degraded by the proteasome or the autophagy/lysosome pathway, alter their subcellular localization, modify their activity or regulate their interaction with other proteins ^2,3^.

βTrCP is considered both an oncoprotein and a tumor suppressor protein, depending on its overexpression or the presence of mutated forms in certain cancers, as well as the substrates with which it interacts and facilitates their ubiquitylation ^4^. In the context of signal transduction, it is important to highlight the role of βTrCP in regulating the WNT/β-catenin, PI3K/AKT/mTOR, ERBB or NF-κB signaling pathways, among others ^5–7^. It also intervenes in the regulation and progression of the cell cycle by degrading various substrates involved in this process, including WEE1, EMI1, CDC25, cyclin D1, cyclin F or PLK1 ^8–11^. Likewise, βTrCP plays various roles in the response to DNA damage by promoting cell cycle arrest, which allows for the processing and repair of the damage. The activation of CHK1 and CHK2 causes the phosphorylation of CDC25, promoting its ubiquitylation by SCF(βTrCP) and subsequent degradation, thereby reducing active cyclin-CDK complexes ^12^. On the other hand, SCF(βTrCP) also promotes the degradation of MDM2, a negative regulator of p53, thus favoring the accumulation of this protein and the consequent cell cycle arrest through the induction of p21 CIP1 ^13^. Additionally, SCF(βTrCP) also plays a role in cell cycle arrest to repair damage following UV radiation by inducing securin degradation ^14^. Finally, once the damage is repaired, SCF(βTrCP) induces the degradation of specific substrates such as WEE1 or claspin, which are involved in the ATR-CHK1 pathway, thereby promoting cell cycle progression ^11,15^. Moreover, βTrCP also regulates substrates involved in the transcription of certain genes, such as PER1, PER2 or FOXO3; in phenomena of cell migration and epithelial-mesenchymal transition, such as SNAIL or TWIST; in the regulation of the apoptotic cascade, such as MCL1 or procaspase 3; in processes related to the immune response, such as IRAK1 or PD-L1; or in viral infections ^4^. In fact, the degradation of the CD4 protein in the context of HIV-1 infection was the first activity described for human βTrCP^16^.

Mammalian cells express two *βTrCP* paralogous genes, *βTrCP1* (also called *FBXW1*) and *βTrCP2* (also called *FBXW11*), which appear to have redundant functions in substrate regulation ^17^, although this may not be universally applicable. In fact, some proteins are specifically degraded by one of the two isoforms. For example, FOXO3 degradation is mediated by βTrCP1, and BST-2 by βTrCP2 ^18,19^. Furthermore, both isoforms can homodimerize or heterodimerize, recognizing substrates in the SCF complex in the first case, and leading to their self-degradation in the second ^20^.

Recently, it has been reported that βTrCP2 inhibits autophagy and senescence, and promotes cell proliferation and migration, while βTrCP1 suppresses cell growth ^20^. In this regard, it has been shown that βTrCP2 preferentially degrades DEPTOR and REDD1, two known inhibitors of mTORC1. This complex promotes cell growth and survival by regulating protein synthesis and other processes, while limiting autophagy-mediated catabolism ^21^. However, mTORC1 complex is constitutively active in senescent cells induced by DNA damage, oncogene expression or replicative stress, even in the absence of amino acids or serum deprivation, preventing full activation of starvation-induced autophagy ^22^. Therefore, βTrCP2 must inhibit senescence through another pathway.

Silencing of *βTrCP2* not only increases the levels of DEPTOR and REDD1, promoting autophagy, but also augments the amount of βTrCP1. We therefore set out to determine the effect of increased amounts of βTrCP1 on cell proliferation and senescence. To this end, we utilized a lentiviral vector and treated the cells with cisplatin as a senescence-inducing agent. Under these conditions, we found that *βTrCP1* overexpression triggers senescent cells’ death through proteasome-dependent degradation of p21 CIP1. Specifically, βTrCP1 disrupts the interaction between p21 CIP1 and nucleophosmin 1 (NPM1), thereby promoting p21 CIP1 degradation, cell cycle progression and cell death. These findings lay the groundwork for targeting βTrCP1/NPM1 interaction as a potential strategy to eliminate senescent cells induced by chemotherapeutic treatment.

## RESULTS

### Overexpression of *βTrCP1* reduces the proliferation rate of hydrogen peroxide-treated cells

As described above, the depletion of *βTrCP2* induces senescence, but the molecular mechanisms underlying this process are not well understood. Since *βTrCP2* depletion also increases βTrCP1 levels, we decided to study the effect of this increase on senescence. To achieve this, we cloned *HA βTrCP1* into a lentiviral vector and infected several cell lines. We confirmed the overexpression of *βTrCP1* by RT-quantitative PCR and verified the functionality of the protein by assessing the degradation of several known SCF(βTrCP1) substrates (Fig. 1A and 2B). We then compared the cell proliferation of the wild-type U2OS cell line with the transduced cell line U2OS::*HA βTrCP1* and found no significant difference between them (Fig. 1C, upper panel). Furthermore, overexpression of *βTrCP1* alone did not induce senescence, as determined by measuring lysosomal β-galactosidase activity, the most widely used biomarker for senescent cells (Fig. 1C, lower panels).

**Figure 1.**
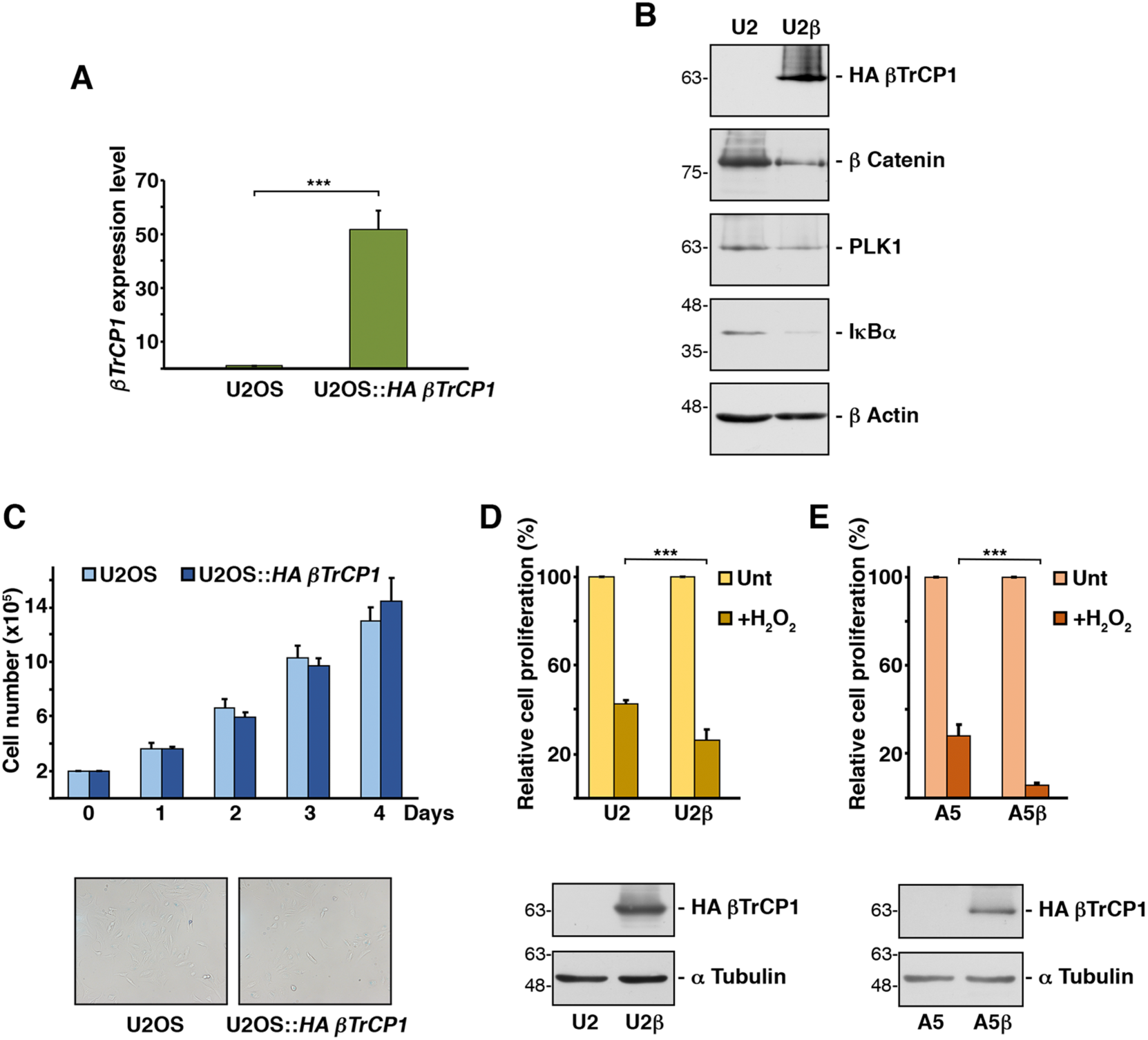
Lentiviral expression of *HA βTrCP1* disminishes the proliferation rate of hydrogen peroxide-treated cells. **A.** Expression levels of *βTrCP1* mRNA in the indicated cell lines. Histograms represent the quantification of mRNA levels by RT-quantitative PCR, using the *GAPDH* as an endogenous control. Error bars represent the standard deviation (SD) (n=3). ***p < 0.001 (Student’s *t*-test). **B.** Whole cell extracts from U2OS (U2) and U2OS::*HA βTrCP1* (U2β) cells were analyzed by Western blotting using the indicated antibodies. **C.** A total of 2x10^5^ cells per cell line were cultured in 10 cm diameter Petri plates, and viable cells were counted daily (upper panel). Error bars represent the SD (n=3). The lower panels show the lysosomal β-galactosidase activity of each cell line using the corresponding kit from Cell Signaling Technology. **D.** Equal numbers of U2OS (U2) and U2OS::*HAβTrCP1* (U2β) cells were cultured in Petri plates and analyzed 3 days later. The protein content in cell extracts was used as a measure of cell proliferation. “+H_2_O_2_” indicates cells treated with H_2_O2 (50 μM) for 2 hours. Unt: untreated cells. Error bars represent the SD (n=3). ***p < 0.001 (Student’s *t*-test). Extracts were also used to analyze HA βTrCP1 protein by Western blotting (lower panels). **E.** Similar to D, but using A549 (A5) and A549::*HA βTrCP1* (A5β) cell lines. Unt: untreated cells.

**Figure 2.**
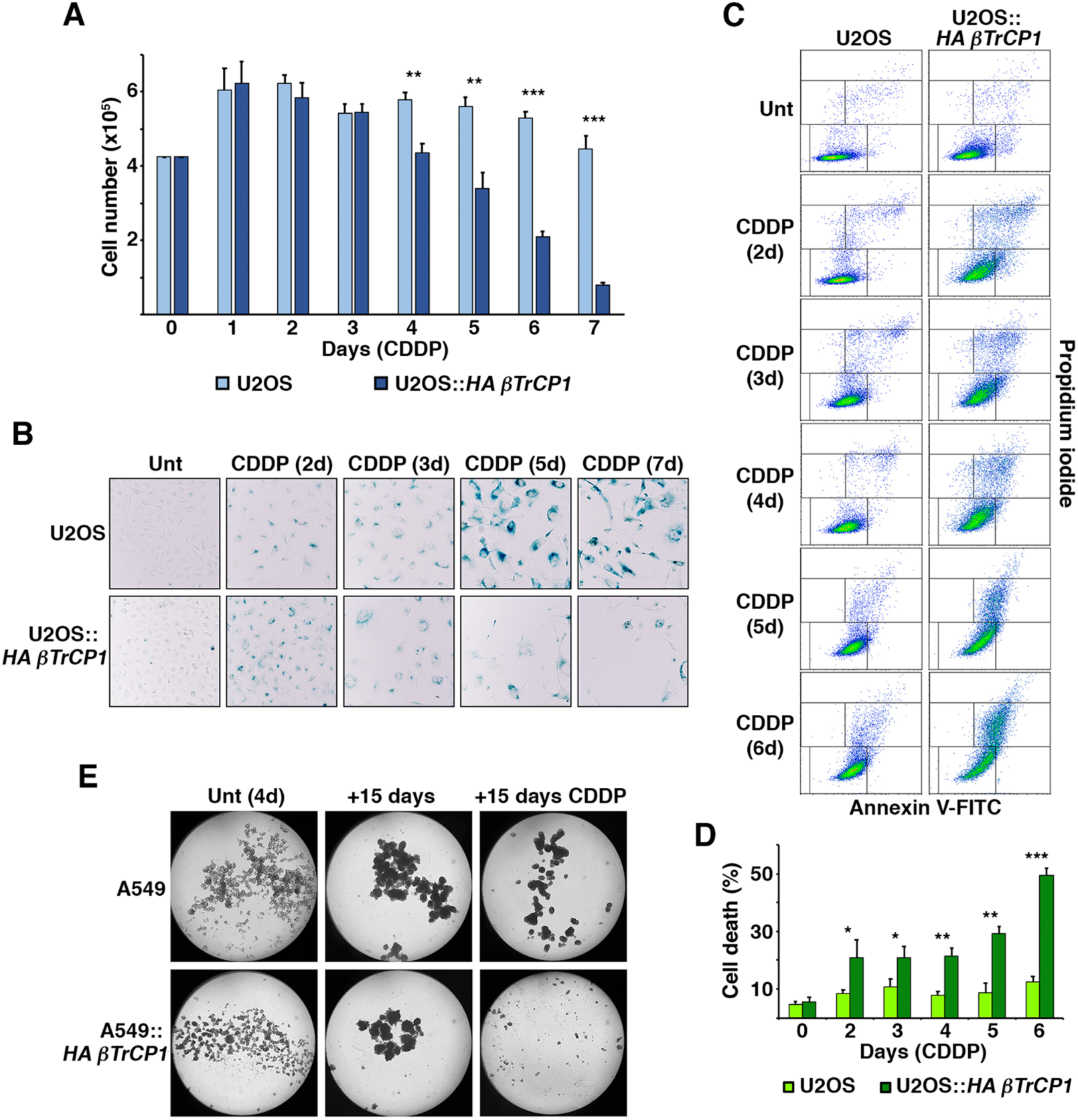
Overexpression of *βTrCP1* induces the elimination of senescent cells derived from cisplatin treatment. **A.** Proliferation of U2OS and U2OS::*HA βTrCP1* cells treated with CDDP (5 μM) was monitored by counting viable cells with a hemocytometer for 7 days. Error bars represent the SD (n=3). **p < 0.01, ***p < 0.001 (Student’s *t*-test). **B.** U2OS and U2OS::*HA βTrCP1* cells were treated with CDDP (5 μM) for the indicated days and assayed for SA-β-gal. Untreated (Unt) cells served as control. **C.** Flow cytometry analysis of annexin V binding assay in U2OS and U2OS::*HA βTrCP1* cells treated with CDDP (5 μM) for the indicated days. Unt: untreated cells. **D.** Percentage of cell death in the annexin V binding experiments described in C. Error bars represent the SD (n=3). *p < 0.05, **p < 0.01, ***p < 0.001 (Student’s *t*-test). **E.** A549 and A549::*HA βTrCP1* cells, cultured under a non-adhesive culture system for 4 days, were treated or not with CDDP (5 μM) for 15 days. The figure shows representative images of tumorspheres obtained through microscopy under each experimental condition.

Next, to further explore the effect of *βTrCP1* overexpression on cells, we subjected them to oxidative stress to induce premature senescence. For this, we used a low concentration of hydrogen peroxide for a short period of time. Specifically, adherent cells were treated for 2 hours with 50 μM H_2_O_2_ and collected 3 days later. As expected, hydrogen peroxide reduced the cell proliferation of U2OS cells to 42.6% of that observed under normal culture conditions (23), likely due to the gradual arrest of cell proliferation and entry into senescence. However, the reduction in cell proliferation in U2OS::*HA βTrCP1* cells treated with hydrogen peroxide was significantly greater than that in U2OS cells (23.6% versus 42.6% in U2OS) (Fig. 1D). To confirm and extend these results, we repeated the experiments in the genetic background of A549 cells. In this case as well, the *βTrCP1*- overexpressing cell line showed a lower number of cells compared to the parental cell line when both were treated with H_2_O_2_ (Fig. 1E). These results indicate that *βTrCP1* overexpression reduces the number of cells when exposed to oxidative stress-induced senescence, either by triggering an early arrest in cell proliferation or leading to cell death.

### Senescence-inducing cisplatin treatment promotes cell death in *βTrCP1*- overexpressing cells

Considering the previous results, we decided to study whether the genotoxic compound cisplatin, which also acts as an oxidative stress-inducing agent and is a standard of care for cancer treatment, would have the same effect. Cisplatin is one of the most widely used chemotherapy drugs for the treatment of solid tumors. However, it has been observed that it can be retained in tissues and organs even after the treatment has been stopped, causing prolonged DNA damage at sublethal doses, which can lead to the so-called therapy- induced senescence ^23^, with possible effects on the maintenance and development of cancer cells.

First, we determined the cisplatin concentration able to induce senescence in our cell models (p16 INK4A-deficient cells) and culture conditions. For this purpose, we cultured U2OS and A549 cells at increasing cisplatin concentrations for 3 days and analyzed the accumulation of p21 CIP1 and p53 by Western blot, as well as the cellular morphological changes characteristic of senescence. The increase in cell size and flattening began at 2.5 μM CDDP, was complete at 5 μM, and cell death started at 10 μM. We chose the concentration of 5 μM because it corresponded to the peak expression of p21 CIP1 along with moderate expression of p53 (Suppl Fig. S1A). To confirm the suitability of this concentration, we measured lysosomal β-galactosidase activity observing a clear blue labeling in both cell models (Suppl Fig. S1B). Next, we analyzed the effect of *βTrCP1* overexpression by comparing the proliferation of U2OS *vs.* U2OS::*HA βTrCP1* cells cultured in the presence of 5 μM CDDP. Figure 2A shows that the number of U2OS::*HA βTrCP1* cells decreased significantly from the fourth day of treatment compared to the parental cell line. Furthermore, from the second day, cells showed lysosomal β- galactosidase activity, indicating that the reduction in cell number occurred once they had reached senescence (Fig. 2B). Similar results were obtained in the A549 cell model (data not shown). To confirm that cell death occurred in cell cultures overexpressing *βTrCP1*, we analyzed the presence of annexin V by flow cytometry. Figures 2C and 2D show the appearance of U2OS::*HA βTrCP1* cells positive for annexin V when treated with cisplatin, which was especially significant from the fourth day of treatment onwards. These data indicate that induction of senescence with cisplatin in cells overexpressing *βTrCP1* causes cell death. Finally, we asked whether this effect could also be observed in three-dimensional (3D) cell cultures, which more closely resemble the natural cellular environment. To this end, we cultured A549 cells in a non-adherent culture system to generate tumorspheres (see Material and Methods). After confirming that they expressed stem cell markers (Suppl Fig. S2), tumorspheres from A549 and A549::*HA βTrCP1* cells were cultured in the presence of 5 μM CDDP. 15 days later, the spheres formed by cells overexpressing *βTrCP1* had practically disappeared, while A549 spheres remained similar to those obtained without cisplatin. Taken together, these results demonstrate that overexpression of *βTrCP1* elicits cell death of cisplatin-induced senescent cells, both in 2D and 3D cultures.

### *βTrCP1*-overexpressing cells show reduced levels of p21 CIP1 following cisplatin-induced senescence

Next, we explored the molecular mechanism by which βTrCP1 induces the death of senescent cells. It has been reported that in p16 INK4A-deficient cells, DNA damage-induced senescence is completely dependent on p21 CIP1, both for its induction and maintenance. In fact, p21 CIP1 does not decrease during the process ^24^. Therefore, we decided to analyze the levels of p21 CIP1 in U2OS and U2OS::*HA βTrCP1* cells treated with cisplatin (5 μM) for 7 days. We observed that, indeed, in U2OS cells p21 CIP1 was induced and remained constant throughout the experiment. However, in U2OS::*HA βTrCP1* cells, the level of p21 CIP1 did not reach that of U2OS cells, and from the fourth day of treatment, it progressively decreased (Fig. 3A and 3B). Similar data were obtained in the A549 cell model (Suppl Fig. S3A and S3B).

**Figure 3.**
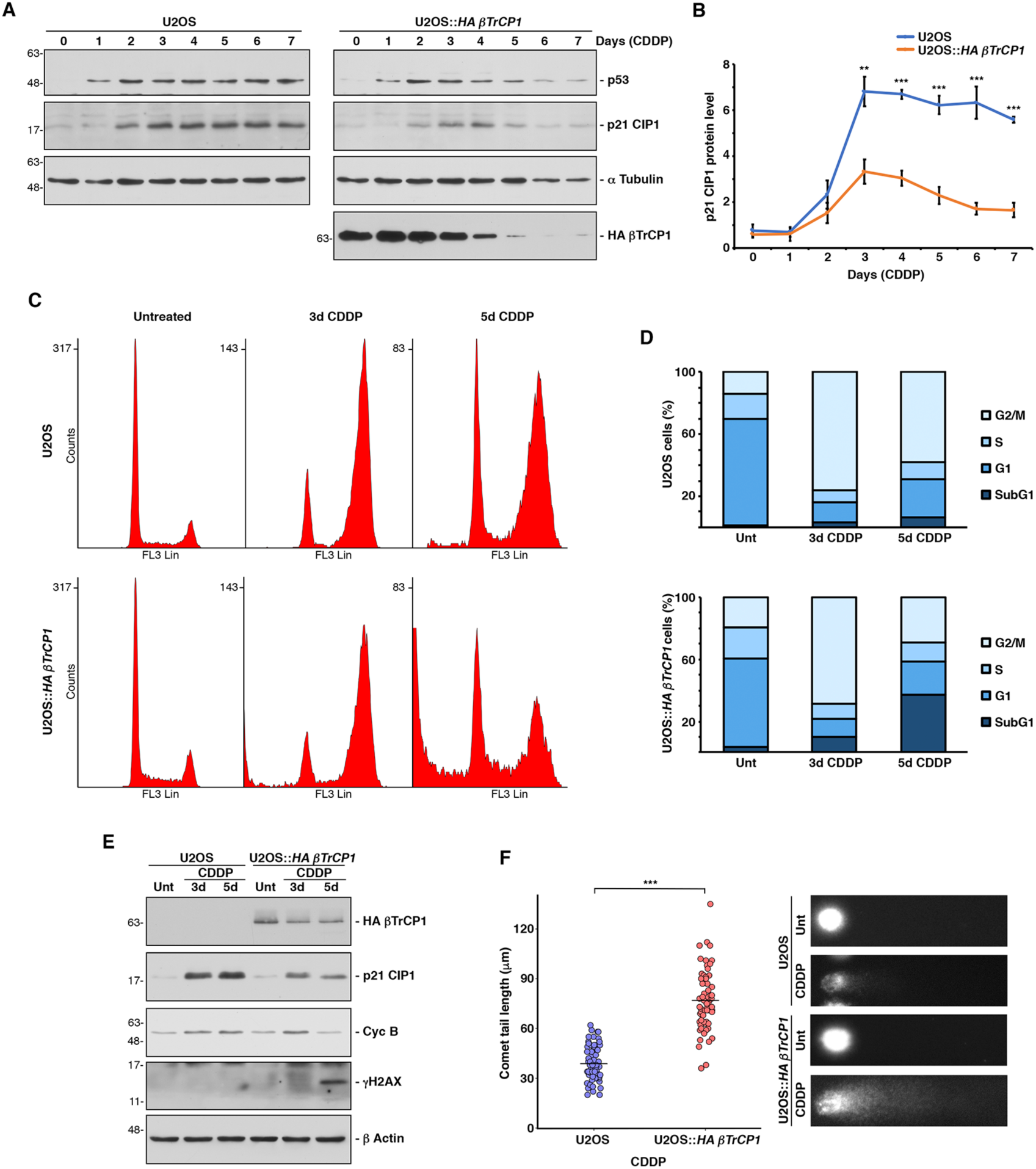
Overexpression of *βTrCP1* in cells cultured under cisplatin-induced senescence conditions reduces p21 CIP1 levels, thereby allowing cell cycle progression and leading to cell death. **A.** U2OS and U2OS::*HA βTrCP1* cells were cultured with CDDP (5 μM), and whole cell extracts were analyzed by Western blotting with the indicated antibodies. **B.** The graph shows the quantification of p21 CIP1 levels using ImageJ software from 3 independent experiments equivalent to that shown in A. Error bars represent the SD. **p < 0.01, ***p < 0.001 (Student’s *t*-test). **C.** Cell cycle analysis by flow cytometry of cells treated with cisplatin or not for 3 and 5 days and subsequently stained with propidium iodide. **D.** Percentage of cells in each phase of the cell cycle from experiments shown in C. Unt: untreated cells. **E.** Western blot of extracts from cells used in C. **F.** U2OS and U2OS::*HA βTrCP1* cells were cultured with CDDP (5 μM) for 5 days and then tested for DNA damage using the comet assay. Each dot represents an individual cell’s comet-tail length (at least 50 cells). The line represents the average tail length of the cells for each cell line. ***p < 0.001 (Student’s *t*-test). Representative DNA tails are shown on the right. Unt: untreated cells.

To determine how the decrease in p21 CIP1 affects the cell cycle arrest induced by this cyclin-dependent kinase inhibitor, we analyzed the cell cycle using flow cytometry and examined cell extracts by Western blot. Our results showed that, under our experimental conditions, U2OS cells arrest in G2/M phase after treatment with cisplatin, while this arrest is less effective in U2OS::*HA βTrCP1* cells. Specifically, after 5 days of treatment, U2OS::*HA βTrCP1* cells exhibited a significant decrease in the G2/M arrested population that corresponds to an increase in the sub-G1 population (Fig. 3C and 3D). This progression through the cell cycle, accompanied by cell death, was corroborated by analyzing the levels of Cyclin B, a cell cycle marker, and γH2AX, a marker of apoptotic DNA fragmentation (Fig. 3E, lane 6 compared to lane 3). Finally, to assess DNA damage at the single-cell level caused by cell cycle progression in the presence of senescence-inducing cisplatin, we performed a single-cell gel electrophoresis assay. Figure 3F shows a marked increase in DNA damage in U2OS::*HA βTrCP1* cells compared to U2OS cells when both were treated with cisplatin (5 μM). Consequently, cisplatin-induced senescence (CIS) in *βTrCP1*-overexpressing cells leads to a gradual reduction of p21 CIP1, enabling cell cycle progression and increasing DNA damage, which ultimately results in cell death.

### βTrCP1 induces proteasome-dependent degradation of p21 CIP1 in cisplatin-induced senescent cells

Expression of *CDKN1A*, which encodes p21 CIP1, has been shown to be upregulated by p53 in response to DNA-damaging agents ^25^. To determine whether βTrCP1 caused the decrease of p21 CIP1 through p53, we analyzed the levels of this protein in RNA interference assays. The specific depletion of *HA βTrCP1* prevented the decrease of p21 CIP1 after CIS, as expected, but did not substantially affect the levels of p53 (Fig. 4A and 4B). This indicates that the effect of *βTrCP1* overexpression on p21 CIP1 is not mediated by p53. To rule out any other possible transcriptional effect, we analyzed the expression of *CDKN1A* after CIS by RT-quantitative PCR. Figure 4C shows that the relative expression levels of *CDKN1A* induced by cisplatin were similar in both the wild-type line and the line overexpressing *βTrCP1*, indicating that the decrease in p21 CIP1 protein in the latter is not due to alterations in *CDKN1A* transcription.

**Figure 4.**
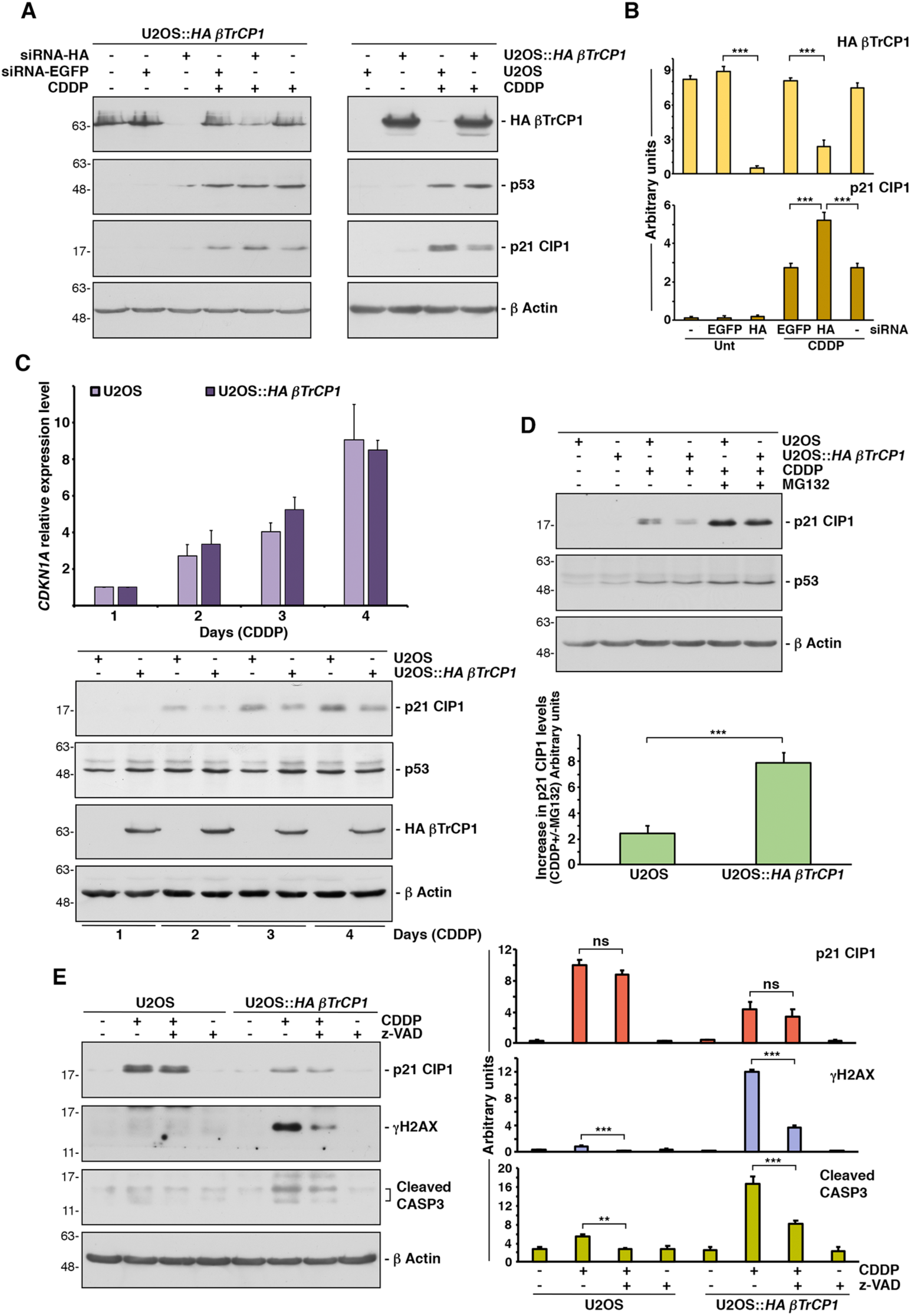
Overexpression of *βTrCP1* triggers the degradation of p21 CIP1 by the proteasome in cisplatin-induced senescent cells. **A.** U2OS::*HA βTrCP1* cells were transfected with the indicated siRNAs and then treated or not treated with CDDP (5 μM) for 3 days (left panels). Whole cell extracts were analyzed by Western blotting using different antibodies. In the right panels, extracts from U2OS and U2OS::*HA βTrCP1* cells were probed with the same antibodies. **B.** Histograms show the quantification of HA βTrCP1 and p21 CIP1 protein levels obtained in A using ImageJ software. Error bars represent the SD (n=3). ***p < 0.001 (Student’s *t*-test). **C.** *CDKN1A* mRNA expression levels in cells treated with CDDP (5 μM) for the indicated days (upper graph). Histograms represent the relative quantification of *CDKN1A* mRNA levels by quantitative RT-PCR, compared to the levels obtained after one day of treatment with 5 μM CDDP. Expression levels were previously normalized to *GAPDH* mRNA levels. Error bars represent the SD (n=3). The relative expression levels of *CDKN1A* did not differ significantly (p > 0.05) (Student’s *t*-test) between the two cell lines throughout the experiment. Cell extracts from the same experiments were used to analyze p21 CIP1 protein by Western blot (lower panels). **D.** Cells treated with CDDP (5 μM) for 3 days were incubated with MG132 3 hours before harvesting. Extracts were analyzed by Western blotting (left panels). The histogram represents the increase in p21 CIP1 levels in cisplatin-treated cells with or without adding MG132. Error bars represent the SD (n=3). ***p < 0.001 (Student’s *t*-test). **E.** Cells were treated with CDDP (5 μM), z-VAD- fmk (20 μM), or both for 3 days. The effect of the treatment on the proteins of interest was analyzed by Western blotting (left panels). Histograms show the quantification of protein levels in left panels using ImageJ software. Error bars represent the SD (n=3). **p < 0.01, ***p < 0.001, ns: not significant (p > 0.05) (Student’s *t*-test).

Next, we used proteasome (MG132) and lysosome (ammonium chloride) inhibitors to determine whether the decrease in p21 CIP1 after CIS was due to a reduction in protein stability. We found that a 24-hour proteasome inhibition prior to cell harvesting nearly completely prevented the reduction of p21 CIP1 levels, in contrast to the effect observed with the lysosome inhibitor (Suppl Fig. S4A and S4B). To avoid the possible toxic effect of the inhibitor, we reduced the treatment time with MG132 to 3 hours, obtaining an equivalent accumulation of p21 CIP1 (Fig. 4D and Suppl Fig. S4C). Additionally, since overexpression of *βTrCP1* induces cell death after CIS, we wanted to rule out a possible role of caspases in the reduction of p21 CIP1. We employed the pan-caspase inhibitor zVAD-fmk, which prevented the accumulation of γH2AX and decreased caspase 3 cleavage, proving its effectiveness, but it did not reverse the decrease in p21 CIP1 after CIS (Fig. 4E). These results together demonstrate that βTrCP1 is responsible for the decrease in p21 CIP1 levels by inducing its degradation via the proteasome.

### Nucleophosmin 1 mediates the effect of βTrCP1 on p21 CIP1

To further investigate the mechanism by which βTrCP1 induces p21 CIP1 degradation, and thereby promotes cell death following CIS, we analyzed the potential interaction between βTrCP1 and p21 CIP1. However, despite multiple attempts, we were unable to detect the *in vivo* interaction between both proteins, if it exists, using co-immunoprecipitation assays. We then proceeded to identify other proteins that might be involved in this relationship. To this end, HA βTrCP1 was immunoprecipitated using both anti-HA and anti-βTrCP antibodies from extracts of U2OS::*HA βTrCP1* cells, treated or untreated with cisplatin (5 μM) for 2 days, with MG132 added 3 hours before harvesting. The immunoprecipitated proteins were identified by mass spectrometry (Suppl Table S1). Among the identified proteins was nucleophosmin 1 (NPM1), which was predicted by UbiBrowser 2.0 database ^26^ as a potential substrate for βTrCP1. NPM1 is a nucleolar protein that translocates to the nucleoplasm under certain conditions, such as after treatment with DNA-damaging agents ^27^. Moreover, in 50% of acute myeloid leukemias, NPM1 is mutated and exhibits cytoplasmic localization ^28^. Interestingly, among its various functions, NPM1 interacts with p21 CIP1 and prevents its ubiquitylation and degradation, thereby prolonging its half-life ^29^.

To investigate the relationship between βTrCP1, NPM1, and p21 CIP1, we first confirmed the βTrCP1/NPM1 association through co-immunoprecipitation assays. For this purpose, cells were treated or not with cisplatin (5 μM), and two fractions were prepared: C fraction, containing soluble proteins extracted with NP40 buffer, and N fraction, containing the remaining proteins, including those associated with chromatin. As expected, NPM1 was found in the latter fraction (Suppl Fig. S5A). Immunoprecipitation of HA βTrCP1 from U2OS::*HA βTrCP1* cells under these conditions revealed that HA βTrCP1 associates with NPM1 only in N fraction, particularly under control conditions (Fig. 5A). In other words, treatment with cisplatin reduced the interaction between these two molecules. This result was confirmed in the wild-type cell line, where the interaction after cisplatin treatment was minimal (Fig. 5B), suggesting that in the U2OS::*HA βTrCP1* cell line, the interaction is detected under these conditions due to the overexpression of *HA βTrCP1*. The complementary experiment, in which NPM1 was immunoprecipitated from the N fraction, confirmed its interaction with HA βTrCP1, as well as a decreased interaction following cisplatin treatment (Fig. 5C).

**Figure 5.**
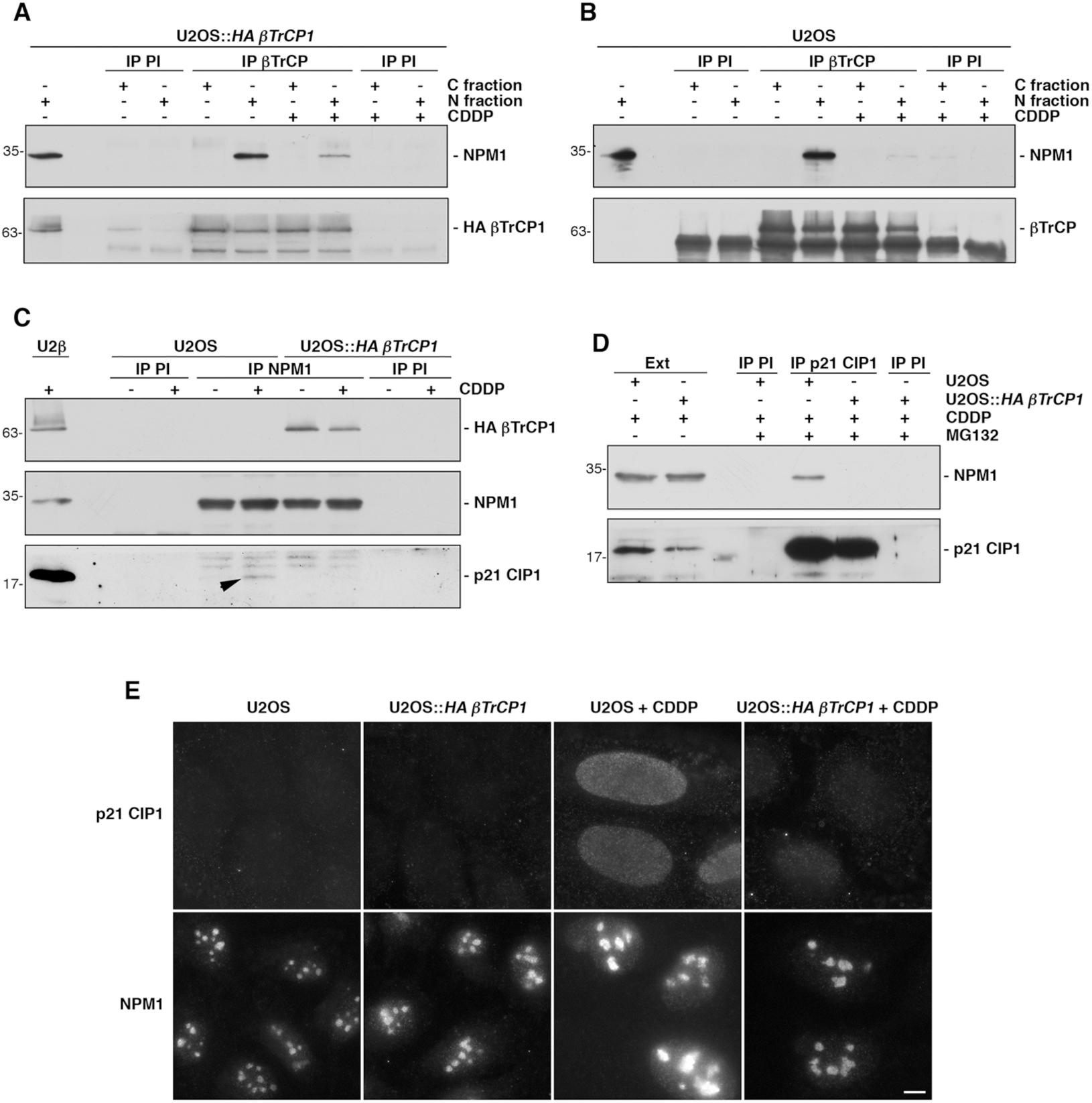
The interaction of HA βTrCP1 with NMP1 prevents the NPM1/p21 CIP1 association, facilitating the degradation of p21 CIP1. **A.** Extracts from C and N fractions of U2OS::*HA βTrCP1* cells, treated or not with CDDP (5 μM) for 3 days, were used to immunoprecipitate βTrCP. Normal mouse serum (PI) was used as an IgG control. Immunoprecipitated materials were analyzed by Western blotting. **B.** Similar experiment to that in A was conducted using U2OS cells. **C.** The N fraction of U2OS and U2OS::*HA βTrCP1* (U2β) cells, untreated or treated with CDDP (5 μM) for 3 days, was used to immunoprecipitate NPM1. Normal mouse serum was also used as a control. Western blots were developed with the indicated antibodies. The arrowhead indicates the co-immunoprecipitation of p21 CIP1 (lane 5). **D.** N fractions were prepared from U2OS and U2OS::*HA βTrCP1* cells treated with CDDP (5 μM) for 3 days, and with MG132 added 3 hours before harvesting. Immunoprecipitations were performed with an anti-p21 CIP1 mouse monoclonal antibody or with normal mouse serum, and the complexes were immunoblotted with the indicated antibodies. Ext: extracts from cells before the addition of the MG132 inhibitor. **E.** U2OS and U2OS::*HA βTrCP1* cells treated with CDDP (5 μM) for 3 days were analyzed by light microscopy. Note the increase in NPM1 in the nucleoplasm of U2OS + CDDP compared to that observed in U2OS::*HA βTrCP1* + CDDP. The bar represents 10 μm.

In the same figure, we can also observe the interaction between NPM1 and p21 CIP1, an association that is lost in cells overexpressing *βTrCP1* (lane 5 *vs.* lane 7), which could ultimately be responsible for the decrease in p21 CIP1 levels following cisplatin treatment and the consequent death of senescent cells. Indeed, downregulation of *NPM1* by siRNA leads to a decrease in p21 CIP1 levels (Suppl. Fig. S5B), as previously described ^29^. Since p21 CIP1 levels are reduced in U2OS::*HA βTrCP1* cells after CIS compared to U2OS cells, we repeated the experiments treating the cells with MG132 before harvesting in order to balance the level of this protein between both cell lines. To avoid the loss of potential p21 CIP1/NPM1 complexes, we performed the assays by immunoprecipitating p21 CIP1, a protein that is much less abundant than NPM1. Figure 5D shows that, despite immunoprecipitating similar amounts of p21 CIP1, it does not interact with NPM1 in U2OS::*HA βTrCP1* cells treated with cisplatin, corroborating the results obtained previously. These data indicate that overexpression of *βTrCP1* prevents the p21 CIP1/NPM1 association following CIS. However, after multiple unsuccessful experiments, we were unable to establish that overexpression of *βTrCP1* caused NPM1 degradation (Suppl. Fig. S5C). Therefore, we investigated whether overexpression of *βTrCP1* could alter NPM1 localization. To this end, we performed microscopy assays and determined that, as expected, NPM1 partially translocates to the nucleoplasm after cisplatin treatment in the wild-type cell line. However, in cells overexpressing *βTrCP1*, NPM1 is retained in the nucleoli (Fig. 5E). Thus, we conclude that overexpression of *βTrCP1* prevents the partial translocation of NPM1 to the nucleoplasm, thereby inhibiting its association with p21 CIP1 and leaving it unprotected against ubiquitylation and degradation. This reduction in p21 CIP1 levels allows the cell cycle to improperly progress, leading to the accumulation of DNA damage and cell death.

## DISCUSSION

Cellular senescence is a state of proliferative arrest that prevents the multiplication of cells subjected to different stresses ^30^. Agents able to induce senescence include DNA-damaging chemotherapeutic drugs as cisplatin. This compound is one of the fundamental pillars in the treatment of various types of cancer ^31^. In this context, as senescence is generally considered to be irreversible, its induction could be beneficial for the patient because it would prevent the proliferation of tumor cells. However, this phenomenon could also be counterproductive for two main reasons. First, cells may sometimes escape this state and proliferate again and, second, senescent cells secrete certain bioactive molecules that can stimulate cell proliferation, evasion of apoptosis, or invasive capacity of surrounding cells ^32^. Therefore, cancer treatments must include strategies both to prevent the emergence of resistant cells and to eliminate senescent cells, avoiding the potential future progression of tumor cells.

Mechanistically, senescence is primarily mediated by the activation of the p53/p21 CIP1 and p16 INK4A pathways, which inhibit CDKs, activate the tumor suppressor RB1 and ultimately lead to cell cycle arrest ^33^. However, in the specific context of DNA damage-induced senescence, whether triggered by adduct-forming agents or oxidative damage inducers, p21 CIP1 alone has been shown to effectively halt cell proliferation ^24^. In this scenario, p21 CIP1 inhibits CDK1, CDK2 and CDK4 during the induction phase of senescence, as well as CDK4 during the maintenance phase. In fact, in our cellular models (U2OS osteosarcoma cells and A549 lung adenocarcinoma cells), both of which are deficient in p16 INK4A, the cells maintained a proper state of senescence after treatment with cisplatin along the days, even reaching truly long-lasting periods of time.

In the present study, we investigated the effect of *βTrCP1* overexpression on cell proliferation and senescence. We observed that increased levels of βTrCP1 do not affect neither cell viability nor proliferation rate when cells are cultured under standard conditions, likely because its activity is tightly regulated ^17^. In contrast, after inducing senescence with cisplatin, cells that overexpress *βTrCP1* ultimately die, and this occurs in both 2D and 3D cultures, which better replicate the natural conditions of tissues or microtumors ^34^. Therefore, although the depletion of *βTrCP2* leads to an increase in βTrCP1 ^20^, this is not responsible for the senescence induced under these conditions.

As we have shown, the death of cells overexpressing *βTrCP1* after CIS is due, at least in part, to the retention of the chaperone NPM1 in the nucleoli, caused by an improper and excessive interaction with βTrCP1. Under senescence conditions, NPM1 is expected to relocate to the nucleoplasm, where it associates with p21 CIP1 and protects it from ubiquitylation and subsequent degradation by the proteasome ^29^. However, in U2OS::*HA βTrCP1* cells after CIS, the p21 CIP1/NPM1 interaction is not detected, leading to the degradation of p21 CIP1, cell cycle progression, consequent accumulation of DNA damage, and ultimately cell death. Similar conclusions can be drawn from previous results showing that *NPM1* depletion reduces p21 CIP1 levels. Nevertheless, if cells are treated with actinomycin D, which causes a redistribution of NPM1 from the nucleoli to the nucleoplasm, the small amount of NPM1 remaining in the cells moves to the nucleoplasm and protects p21 CIP1, increasing its levels in the cells ^35^.

NPM1 is a multifunctional protein involved in various processes such as ribosomal maturation, centrosome replication, maintenance of genomic stability, DNA repair, cell cycle control and apoptosis ^36^. Among its numerous partners, NPM1 interacts with p14 ARF, localizing it to the nucleolus in non-stressed cells. After different stimuli, p14 ARF translocates to the nucleoplasm and interacts with MDM2, facilitating the accumulation of p53 ^37^. A direct interaction between NPM1 and MDM2 has also been demonstrated, protecting p53 from degradation as a result ^38^, as well as an interaction with p53 itself ^39^, although some researchers question this ^40^, or with p21 CIP1 ^29^, as we have previously described. The extensive interaction network of NPM1 complicates the understanding of its role in cellular functions. At present, we do not know how the excess of βTrCP1 retains NPM1 in the nucleolus. Since this protein lacks a canonical nucleolar localization signal, it is believed that its retention in the nucleolus is due to its association with other resident proteins and rRNA ^41^. Upon analyzing various agents that induce cellular stress, it has been shown that all of them lead to nucleolar oxidation, which triggers the translocation of NPM1 through S-glutathionylation of cysteine 275 ^42^. Under these conditions, the βTrCP1/NPM1 interaction is significantly disrupted, but the excess βTrCP1 manages to partially maintain it, which may lead to NPM1 nucleolar retention. However, further studies are needed to clarify this mechanism more precisely.

Other authors have also reported alterations in the translocation of NPM1 to the nucleoplasm and how this affects senescence. In the endothelial cell model, where senescence plays a key role in the onset and progression of cardiovascular diseases, overexpression of nuclear Heme oxygenase-1 (HO-1) prevented the translocation of NPM1 from the nucleolus to the nucleoplasm, inhibiting p53 activation and, consequently, senescence ^43^. From a therapeutic perspective, it remains unclear whether preventing senescence under these stress conditions is beneficial. In any case, when senescence is induced by sublethal doses of DNA-damaging agents, as may occur along anti-tumor treatments, eliminating those cells potentially altered by chemotherapeutics is likely more critical. In fact, entry into senescence can be considered a mechanism of resistance to chemotherapy drugs. Cisplatin, for example, often elicits an initially successful therapeutic response. However, apart from potential side effects, certain tumors are intrinsically resistant to the drug, while others that are initially sensitive gradually develop resistance ^44^. One of the most notable examples is melanoma.

Traditional therapies, such as cisplatin treatment, are largely ineffective. It has recently been discovered that one of the underlying causes is that melanoma cells are particularly prone to entering senescence ^45^. Furthermore, in these cases, the senescence-associated secretory phenotype is particularly robust, inducing a high rate of proliferation in neighboring non-senescent cells. Therefore, both in these cases, and in others where senescence is acquired after treatment has concluded, it becomes crucial to eliminate senescent cells.

Senolytic drugs are small molecules designed to selectively eliminate senescent cells. However, these cells are not easy to kill, partly because they are protected from apoptosis by upregulating certain proteins involved in anti-apoptotic pathways ^46^. So far, a limited number of senolytic targets have been identified, including BCL-XL, HSP90, BRD4, OXR1, and FOXO4 ^47^. However, current senolytics still present some limitations in terms of safety and specificity. Therefore, identifying new senolytic targets is crucial for the design and development of more effective senolytics. Whether NPM1 can be a target for senolytic drugs remains to be determined. Nonetheless, we know that if we manage to retain this molecule in the nucleolus after inducing senescence with DNA-damaging agents, these senescent cells ultimately die.

## MATERIALS AND METHODS

### Plasmids and viral transductions

pLenti 6.3 HA βTrCP1, previously described ^48^, along with pMD2-G, pMDLg-pRRE and pRSV-Rev from Addgene (Watertown, MA, USA), were transfected into HEK293T cells using Xfect reagent (Clontech, Mountain View, CA, USA). After 48 hours of transfection, the cell supernatant was harvested, filtered through a 0.45 μm Puradisc 25 mm filter (GE Healthcare, Little Chalfont, UK), and the viruses were concentrated using a Lenti-X concentrator (Clontech). U2OS and A549 cells were transduced for 24 hours and selected after seven days in 20 μg/mL blasticidin (Gibco, Thermo Fisher Scientific, Waltham, MA, USA).

### Cell culture, transfections, drugs, and cell lysis

Routinely, U2OS, A549, and HEK293T cells (from ATCC), and their derivatives were grown in Dulbecco’s Modified Eagle Medium (DMEM) from BioWest (Nuaillé, France), as previously described ^49^. To suppress gene expression, cells were transfected with siRNA-NPM1 ^39^ or siRNA-HA (5’-CGC UUA UCC UUA UGA CGU A [dT] [dT]-3’) using the Xfect™ RNA transfection reagent (Takara Bio Inc., Shiga, Japan) following the manufacturer’s standard instructions. siRNA-EGFP ^50^ was used as a non-specific control. Cells were harvested 48 hours after transfection, and the reduction in protein levels was confirmed by Western blotting. In the indicated experiments, cells were treated with cisplatin (CDDP, 2.5-40 μM, Sigma-Aldrich, St. Louis, MO, USA), MG132 (20 μM, Santa Cruz Biotechnology, Dallas, TX, USA), ammonium chloride (40 mM, Sigma-Aldrich), z-VAD-fmk (20 μM, Sigma-Aldrich) or H_2_O_2_ (50 μM, Sigma-Aldrich), and harvested at various time points.

Whole cell extracts were prepared at room temperature (RT) in UTB buffer (8 M urea, 50 mM Tris-HCl pH 7.5, 150 mM β-mercaptoethanol) for 10 min and sonicated. The extracts were heated 5 minutes at 95 °C and centrifuged at 20,000 *g* for 20 min. The supernatants were stored at -80 °C. Protein concentration was determined using the Bradford assay (Bio-Rad, Hercules, CA, USA).

### Electrophoresis, Western blot analysis, and antibodies

Proteins were separated by SDS-polyacrylamide gel electrophoresis (SDS-PAGE) and transferred to nitrocellulose membranes (GE Healthcare). The membranes were then probed with the following antibodies: anti-HA-peroxidase monoclonal antibody (#12013819001, Roche, Basel, Switzerland); anti-β actin (sc-47778 HRP) and anti-p53 (sc-126) monoclonal antibodies, as well as anti-Oct3/4 (sc-8628), anti-Sox2 (sc-17320) and anti-Sox9 (sc-20095) polyclonal antibodies (Santa Cruz Biotechnology); anti-α tubulin monoclonal antibody (T9026, Sigma-Aldrich); anti-cyclin B (#610220) and anti-β catenin (#610154) monoclonal antibodies (BD Biosciences, Franklin Lakes, NJ, USA); anti-PLK1 (#05-844), anti-γH2AX (#05-636) and anti-active β catenin (#05-665) monoclonal antibodies (EMD Millipore, Merck, Darmstadt, Germany); anti-βTrCP (#4394) and anti-cleaved caspase-3 (#9664) monoclonal antibodies, as well as anti-IκBα (#9242) polyclonal antibody (Cell Signaling Technology, Danvers, MA, USA); anti-histone H1 monoclonal antibody (ab11079, Abcam, Cambridge, UK); and anti-p21 CIP1 (GT1032) and anti-NPM1 (GTX57555) monoclonal antibodies (GeneTex, Irvine, CA, USA). Peroxidase-coupled donkey anti-rabbit IgG (NA934V) and sheep anti-mouse IgG (NA931) were purchased from GE Healthcare. Immunoreactive bands were visualized using the Enhanced Chemiluminescence (ECL) Western blotting system (GE Healthcare).

Protein level quantification was carried out using ImageJ software (Image Processing and Analysis in Java; National Institutes of Health, Bethesda, MD, USA; http://imagej.nih.gov/).

### Subcellular fractionation

Cells were resuspended in NP40 buffer (150 mM NaCl, 10 mM Tris-HCl (pH 7.5), 1% NP40, 10% glycerol, 1 mM PMSF, 1 μg/mL aprotinin, 1 μg/mL pepstatin and 1 μg/mL leupeptin) and incubated on ice for 30 minutes. After centrifugation at 20,000 *g* for 20 minutes, the supernatant was collected as the C fraction. The pellet was resuspended either in UTB buffer, as described above, for Western blot analysis, or in NP40 buffer containing 420 mM NaCl, sonicated, and centrifuged for co-immunoprecipitation experiments. This second supernatant was collected as the N fraction. Protein concentration was determined using the Bradford assay (Bio-Rad).

### Co-immunoprecipitation experiments

1-2 mg of protein from the C and N (diluted to 150 mM NaCl) fractions were incubated with normal mouse (sc-2025) or rabbit (sc-2027) sera (Santa Cruz Biotechnology) for 30 minutes, followed by incubation with protein G or A-Sepharose beads (GE Healthcare), respectively, for 1 hour at 4 °C. After centrifugation, the beads were discarded, and the supernatants were incubated for 2 hours with anti-βTrCP (#4394, Cell Signaling Technology), anti-NPM1 (GTX57555, GeneTex), or anti-p21 CIP1 (GT1032, GeneTex) antibodies, or with normal mouse or rabbit sera, followed by protein G or A-Sepharose beads for 1 hour. The beads were washed, and the bound proteins were solubilized by adding SDS sample buffer and heated at 95 °C for 5 minutes.

### Tandem mass spectrometry (MS/MS)

Samples were digested with modified porcine trypsin (Promega, Madison, WI, USA) at a final trypsin-to-protein ratio of 1:50. Digestion was carried out overnight at 37 °C. The resulting peptide mixtures were subjected to nano-liquid chromatography coupled to MS for protein identification. Peptides were injected onto a C-18 reversed-phase (RP) nano-column and analyzed using a continuous acetonitrile gradient. A flow rate of 200 nL/min was employed to elute peptides from the RP nano-column to an emitter nanospray needle for real-time ionization and peptide fragmentation on an LTQ-Orbitrap mass spectrometer (Thermo Scientific). For protein identification, tandem mass spectra were extracted and charge state deconvoluted by Proteome Discoverer 1.4.0.288 (Thermo Scientific). All MS/MS samples were analyzed with SEQUEST™ (Thermo Scientific).

### Cell proliferation assays

Cells were either left untreated, treated with CDDP (5 μM) throughout the entire experiment, or treated with H_2_O_2_ (50 μM) for 2 hours, and then counted at different time points. Adherent cells were trypsinized, collected, and viable cells (those excluding trypan blue) were counted using a hemocytometer. In parallel, as an additional method for measuring cell proliferation, the total protein content was quantified using the Bradford assay (Bio-Rad).

### RT-quantitative PCR

The SuperScript™ IV CellsDirect™ cDNA Synthesis Kit (Thermo Fisher) was used to obtain cDNA from cell extracts. To perform the qPCR experiments, the LightCycler 480 II thermal cycler (Roche) and the LightCycler 480 SYBR Green I Master kit (Roche) were used. The primers used for the amplification of *βTrCP1* and *CDKN1A*, with *GAPDH* as an internal reference, were: βTrCP1-RT-S: 5’-CAT TGT TTC TGC ATC TGG GGA T-3’; βTrCP1-RT-R: 5’-TCA AAT CGA ATA CAA CGC ACC A-3’; CDKN1A-S: 5’-TGT CAC TGT CTT GTA CCC TTG-3’; CDKN1A-R: 5’-GGC GTT TGG AGT GGT AGA A-3’; GAPDH-S: 5’-ACA CCC ACT CCT CCA CCT TT-3’; GAPDH-R: 5’-TCC ACC ACC CTG TTG CTG TA-3’. Analysis was performed using LightCycler 480 software.

### Senescence associated (SA) β-galactosidase staining

Cells (5x10^4^) were seeded in 35 mm diameter plates (Ibidi GmbH, Gräfelfing, Germany) and treated or not with CDDP (5 μM) for the indicated days. The cells were washed twice with PBS, and SA-β-gal was detected using the Senescence β-Galactosidase Staining Kit (#9860, Cell Signaling Technology) according to the manufacturer’s protocol. Cells were incubated at 37 °C for 18 hours and photographed.

### Annexin V binding assays

U2OS and U2OS::*HA βTrCP1* cells were either left untreated or treated with 5 μM CDDP for the indicated times. One million cells were washed in cold PBS, resuspended in 200 μl of annexin V binding buffer containing 10 μl of propidium iodide and 10 μl of annexin V-FITC (Annexin V-FITC Apoptosis Detection Kit I, BD Biosciences), and incubated for 15 minutes at RT in the dark. After incubation, the cells were diluted in 300 μl of annexin V binding buffer. Fluorescence was measured on a MACSQuant VYB flow cytometer (Miltenyi Biotec, Bergisch Gladbach, Germany) within 1 hour. Cell populations (viable, early apoptotic, and late apoptotic) were identified by measuring fluorescence on the B1 Annex-FITC-A and Y2-PI-A channels. Data were analyzed using MACSQuantify™ software.

### Cell cycle analysis by flow cytometry

For cell cycle analysis, cells were collected after various treatments, washed twice with cold PBS, and fixed overnight in 70% cold ethanol at -20° C. Propidium iodide staining was performed after 1-hour incubation at 37 °C with 0.2 mg/mL RNase A solution (Sigma-Aldrich). The cells were then incubated with a 20 μg/mL propidium iodide solution (Sigma Aldrich) for 15 minutes at 4 °C in the dark. DNA content was measured using CXP software on a Cytomics FC500-MPL flow cytometer (Beckman Coulter, Brea, CA, USA). The percentage of cells in each phase of the cell cycle was quantified using CXP software.

### Tumorsphere formation

A549 and A549::*HA βTrCP1* cells were trypsinized and seeded into 6-well culture plates coated with a thin layer of sterile 1.2% agarose (dissolved in water) at a density of 2×10^4^ cells per well. The cells were incubated for 4 days to allow tumorsphere formation. CDDP (5 μM) was then added to selected wells, and the plates were photographed 15 days later.

### Comet assay

Assays were performed as previously described for alkaline unwinding and electrophoresis conditions ^51^. After 30 minutes in alkaline buffer, gel electrophoresis was carried out at 25 V for 30 minutes at 4 °C. The comet assay was performed on cells exposed to CDDP (5 μM) for 5 days or left untreated. Positive control cells were treated with 20 μM actinomycin D for 1 hour. Positive and negative controls were included with each sample. A minimum of 50 individual cells per sample were analyzed in triplicates. Comet analysis was performed using the ImageJ analysis tool. Statistical analysis was conducted using two-way ANOVA.

### Immunofluorescence microscopy

Cells were grown on coverslips, fixed with 4% paraformaldehyde for 10 minutes on ice, and permeabilized with 0.25% Triton X-100 for 5 minutes at RT. The cells were then incubated with blocking buffer (1% BSA, 0.1% Tween 20 in H_2_O) for 1 hour at RT. Coverslips were incubated with primary antibodies for 1 hour at RT, washed three times with 1% PBS and once with blocking buffer, and incubated with the appropriate fluorescent secondary antibody for 1 hour at RT. The coverslips were subsequently washed three times with 1% PBS and mounted on microscopy slides using mounting reagent with DAPI (Ibidi GmbH). Staining was analyzed using a Leica DMi8 inverted microscope with a Plan Apo 63x oil-immersion objective under consistent laser parameters. All microscope images were analyzed with ImageJ software. The primary antibodies used were anti-p21 CIP1 (GT1032, GeneTex, Irvine, CA, USA) and anti- NPM1 (sc-0412, Santa Cruz Biotechnology) mouse monoclonal antibodies.

### Quantification and statistical analysis

Details regarding the statistical analyses, including the number of replicates and the use of standard deviation (SD), are provided in the Figure Legends. Routinely, statistical analyses were performed with unpaired Student’s *t*-tests. Statistical significance is indicated in the figures as *p < 0.05, **p < 0.01 and ***p < 0.001.

## AUTHOR CONTRIBUTIONS

A.B.-F. and F.R. performed study concept and design. A.B.-F., J.H.-R., M.C.L.-M. and, M.M.-S. provided acquisition, analysis and interpretation of data, and statistical analysis.

C.S. and M.A.J. performed development of methodology. C.S., M.A.J. and F.R. wrote the manuscript with input from all other authors. All authors read and approved the final paper.

## DATA AVAILABILITY

All data generated or analyzed during this study are included in this published article (and its supplementary information files).

