## Supplementary figures for "Overexpression of *βTrCP1* elicits cell death in cisplatin-induced senescent cells"

### SUPPLEMENTARY FIGURES AND TABLE

**Supplementary Figure S1.** Determination of the senescence-inducing cisplatin concentration in U2OS and A549 cells.
**A.** Cells were treated with the indicated concentrations of cisplatin for 3 days. Whole cell extracts were subjected to SDS-PAGE, transferred to membranes by electroblotting, and analyzed using various antibodies. **B.** Lysosomal β-galactosidase activity was detected in cells treated or not with CDDP (5 μM) for 3 days.

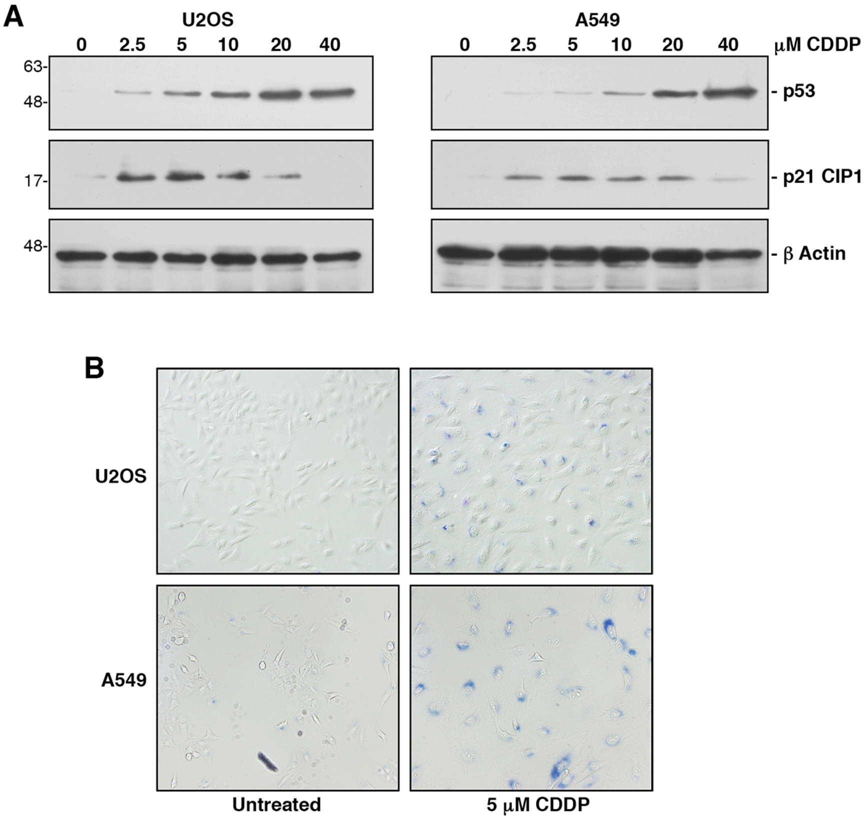

**Supplementary Figure S2.** Expression of stem cell markers in tumorspheres derived from A549 and A549::*HA βTrCP1* cells.
Extracts from cells or tumorspheres, obtained after 10 days of culture under a non-adhesive system, were analyzed by Western blotting using various antibodies.

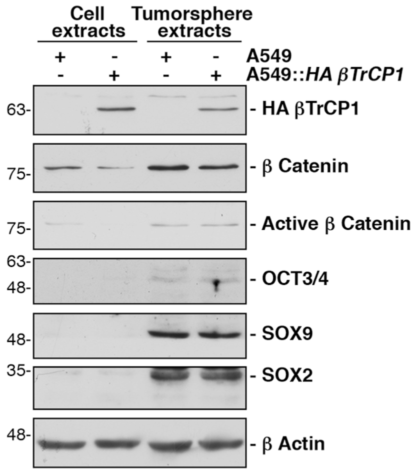

**Supplementary Figure S3.** Overexpression of *βTrCP1* in A549 cells induces a reduction of p21 CIP1 levels in cisplatin-induced senescent cells.

**A.** A549 and A549::*HA βTrCP1* cells were cultured with CDDP (5 µM), and whole cell extracts were analyzed by Western blotting with the indicated antibodies. **B.** The graph shows the quantification of p21 CIP1 levels using ImageJ software from 3 independent experiments equivalent to that shown in A. Error bars represent the SD. **p < 0.01, ***p < 0.001 (Student's *t*-test).

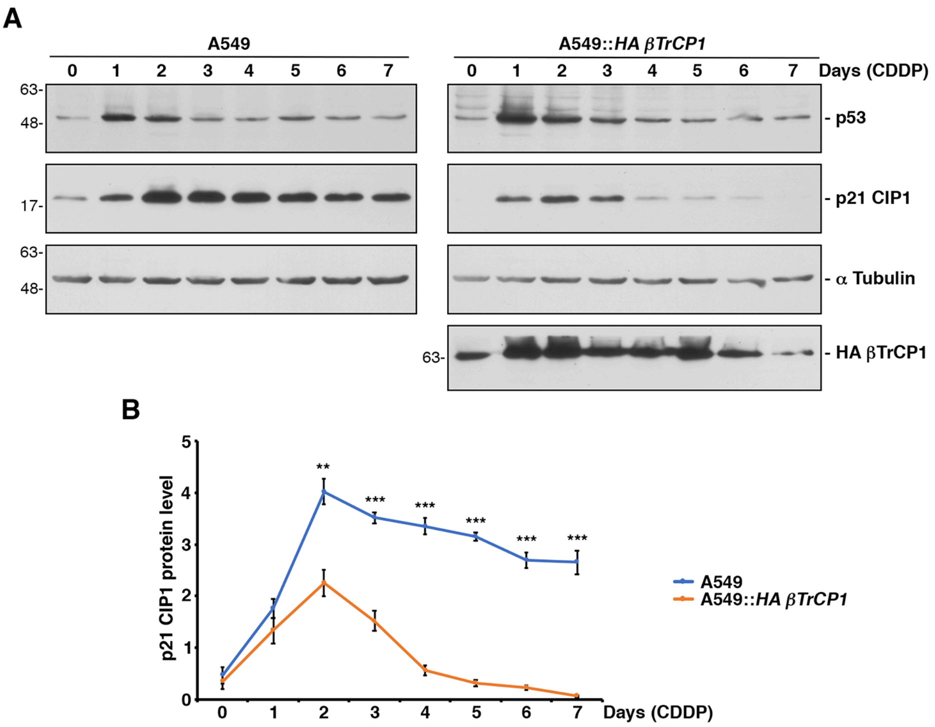

**Supplementary Figure S4.** The overexpression of *βTrCP1* in A549 cells causes degradation of p21 CIP1 by the proteasome in cisplatin-induced senescence conditions. **A.** A549 and A549::*HA βTrCP1* cells were treated with CDDP (5 µM) for 24 (+) or 48 (++) hours. Twenty-four hours before harvesting, cells were incubated with MG132 or $NH_4Cl$. Extracts were analyzed by Western blotting. **B.** The histogram represents the increase in p21 CIP1 levels in cisplatin-treated cells with or without adding MG132. Error bars represent the SD (n=3). ***p < 0.001 (Student's *t*-test). **C.** Similar experiment to A was performed in C, but with cells incubated for 48 hours with CDDP and for 3 hours with MG132 before harvesting (left panels). The histogram represents the increase in p21 CIP1 levels in cisplatin-treated cells with or without adding MG132. Error bars represent the SD (n=3). **p < 0.01 (Student's *t*-test).

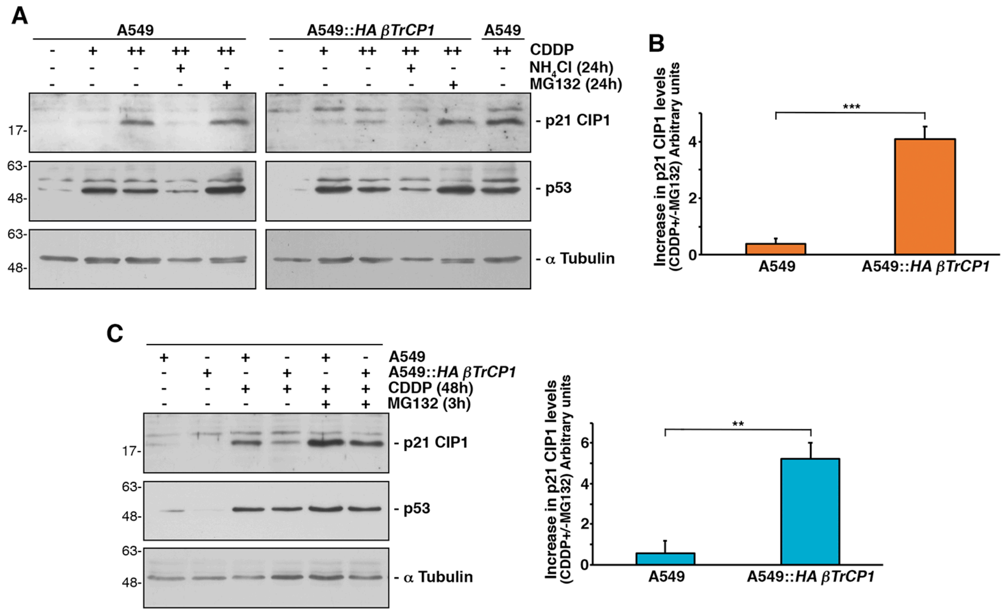

**Supplementary Figure S5.** NPM1, a protein involved in the stability of p21 CIP1, is located in the insoluble fraction of NP40 extracts and is not degraded by the lentiviral expression of *βTrCP1*.
**A.** The subcellular localization of NPM1 was studied in the soluble (C fraction) and insoluble (N fraction) fractions of NP40 extracts from cells treated or not with CDDP (5 μM) for 3 days. The figure shows the corresponding Western blotting. **B.** U2OS cells were transfected with siRNA-NPM1 and treated or not with CDDP (5 μM) for 3 days to determine its effect on p21 CIP1 stability. Whole cell extracts were analyzed using the indicated antibodies. **C.** Extracts from U2OS and U2OS::*HA βTrCP1* cells treated or not with CDDP were subjected to Western blot analysis to investigate the effect of *βTrCP1* overexpression on NPM1 stability.

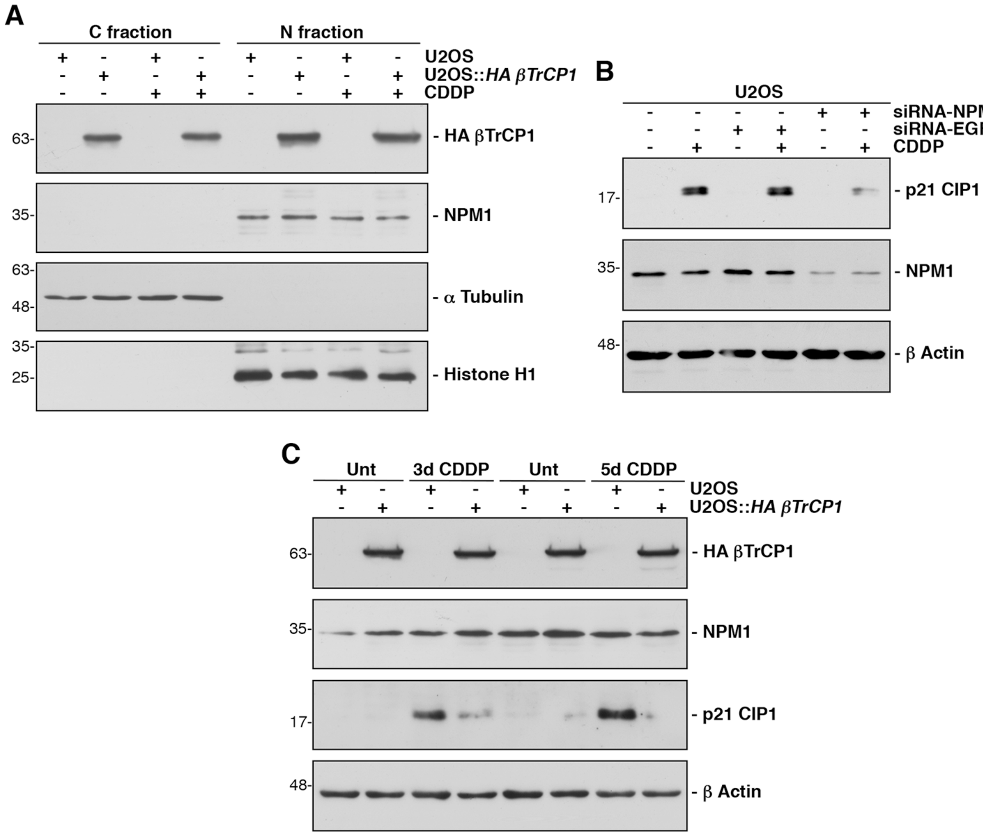

**Supplementary Table S1.** Identification of βTrCP1-interacting proteins by tandem mass spectrometry.

Extracts from U2OS::*HA βTrCP1* cells, treated or not with CDDP (5 μM) for 2 days and with MG132 for 3 hours before harvesting, were used to identify βTrCP1-interacting proteins. Data shown in red correspond to those obtained from the IP HA of control cells. Data shown in green correspond to those obtained from the IP HA of CDDP-treated cells. Data shown in light blue correspond to those obtained from the IP βTrCP of control cells, and data shown in yellow correspond to those obtained from the IP βTrCP of CDDP-treated cells.
